# Cell Type-informed Characterization of Spatial Niches from Spatial Multimodal and Multi-omics Data

**DOI:** 10.64898/2026.05.09.722417

**Authors:** Gaoyuan Du, Jialuo Xu, Xinrong Wei, Chang Liu, Dongmin Zhao, Xiangting Jia, Xingyi Li, Xuequn Shang

## Abstract

Cell niches play critical roles in tissue organization and orchestrate homeostasis, development, and disease progression. Advances in spatial omics technologies now allow diverse molecular and image-derived data to be jointly captured while preserving spatial context, but deciphering cell niches from such spatial multimodal and multi-omics data remains challenging. Existing computational methods are still limited in their flexibility across variable combinations of spatial modalities and omics data. Here we introduce SpaNECT, a unified and flexible framework designed to accommodate spatial multimodal and multi-omics data for cell niche characterization. SpaNECT further incorporates reference-informed cell-type information to support biologically interpretable niche analysis. Systematic evaluations across diverse tissues, disease conditions, and developmental stages showed that SpaNECT consistently outperformed representative methods in resolving cell niches. In mouse brain spatial multi-omics data, SpaNECT uncovered niche-associated molecular and regulatory programs; in developing chick heart, it tracked cross-stage niche reorganization and progressive remodeling of ventricular-associated cell states during maturation. Overall, SpaNECT establishes a general and robust framework for characterizing cell niches across spatial multimodal and multi-omics data.

## 1 Introduction

Cell niches are spatially organized cell communities with coordinated tissue functions and play important roles in tissue homeostasis, development, and disease progression[1–3]. Recent advances in spatial omics technologies, including spatial transcriptomics[4–12] and spatial multi-omics platforms[13–16], now enable the simultaneous acquisition of omics data and histological images while preserving spatial context, providing complementary views for characterizing cell niches in tissues[17]. Despite these opportunities, it remains a challenge to accurately resolve and interpret cell niches from multimodal and multi-omics spatial atlases.

Various computational methods have been developed to identify and characterize cell niches from spatial omics data [18]. Spatial single-omics methods [19–24], such as GraphST[25], stDCL[26], and CytoCommunity[27], commonly use gene expression, spatial coordinates, or local cellular neighborhoods to model tissue organization and cell microenvironments. These methods have advanced spatially informed niche analysis; however, because cell-type composition is a primary feature underlying spatial niches, approaches that mainly rely on transcriptomic heterogeneity may not directly establish the relationship between inferred niches and their constituent cell populations, making cellular interpretation more challenging [27]. Histology-aware methods[28], such as SpaGCN[29], DeepST[30], and STAIG[31], combine spatial gene expression with tissue morphology through morphology-guided graph construction, expression enhancement, or image-aided representation learning. However, in many of these designs, morphology is mainly treated as auxiliary information, often with limited consideration of how image-derived and molecular representations retain their modality-specific cellular context before coordination. Concurrently, spatial multi-omics integration methods[32–35], such as SpatialGlue[36], have advanced the joint modeling of multiple spatial molecular modalities and enabled integrated representations for resolving tissue architecture and molecular programs. However, many existing approaches are developed around specific modality combinations or integration objectives, and their learned representations can be sensitive to variations in modality quality[37]. These developments reveal a broader design challenge. As spatial multimodal and multi-omics studies increasingly involve diverse modality combinations that differ in availability, quality, and information content across datasets [32, 38, 39], less systematic attention has been paid to how such inputs can be coordinated while preserving modality-specific context and enabling biological interpretability of niche representations.

In this study, we introduce SpaNECT, a unified and flexible framework that accommodates diverse combinations of spatial multimodal and multi-omics data for characterizing cell niches. Rather than merging all inputs at the outset, SpaNECT separates the learning of modality-specific cellular context from cross-modal alignment. This design allows each modality to contribute according to its own noise structure and information content, reducing the risk that modality-specific signals are diluted by premature fusion. Moreover, SpaNECT incorporates reference-informed cell-type information to improve biological interpretability by relating learned niche representations to the cell populations that shape local tissue contexts.

Using diverse spatial multimodal and multi-omics datasets from different tissues, disease conditions, and developmental stages [14, 40–43], we benchmarked SpaNECT against representative spatial single-omics and spatial multi-omics methods. In spatial multi-omics data from mouse brain [14], SpaNECT characterized niches through molecular, regulatory, and compositional programs. In developing chick heart sections profiled across developmental stages [43], SpaNECT further tracked cross-stage niche reorganization during maturation.

## 2 Results

### 2.1 Overview of SpaNECT

SpaNECT proceeds in two stages: it first learns modality-specific cellular context on a shared spatial graph and then aligns these contextual representations across modalities to derive a unified niche representation. SpaNECT takes as input feature matrices measured at segmented cells or spatial capture locations, such as spots, pixels, bins, or other spatial units, together with their spatial coordinates. We refer to these spatial units as cells hereafter for brevity and without restricting SpaNECT to a specific technological platform or resolution. By default, SpaNECT uses spatially resolved data from the same tissue section as input, together with spatial coordinates and reference-informed cell-type information derived from available annotations or deconvolution using external single-cell references. These inputs may include spatial transcriptomic data, additional molecular modalities such as proteomic or epigenomic data, and histology-derived image representations when available (Fig. 1a). For each modality, SpaNECT first organizes the input into a modality-specific feature matrix. In parallel, it constructs a shared cellular spatial graph from physical proximity and introduces the cell-type matrix as an explicit component of the model (Fig. 1b). It then applies a Cellular Context-aware Contrastive Encoder (CCCE) to each modality, allowing the resulting embeddings to preserve both local spatial organization and semantic similarity among cells (Fig. 1c). After this context-learning stage, the pre-trained encoders are frozen and complemented with lightweight adapters. The resulting modality embeddings are aligned through a cross-attention module and aggregated into a unified cell-type-informed representation. Modality-aware decoding heads then reconstruct the original modality profiles from this shared representation (Fig. 1b). Finally, SpaNECT provides a suite of downstream analyses for characterizing and comparing cellular niches across biological conditions or developmental stages (Fig. 1d). In addition, SpaNECT can readily incorporate additional modalities without altering the core context-learning and alignment scheme.

**Fig. 1.**
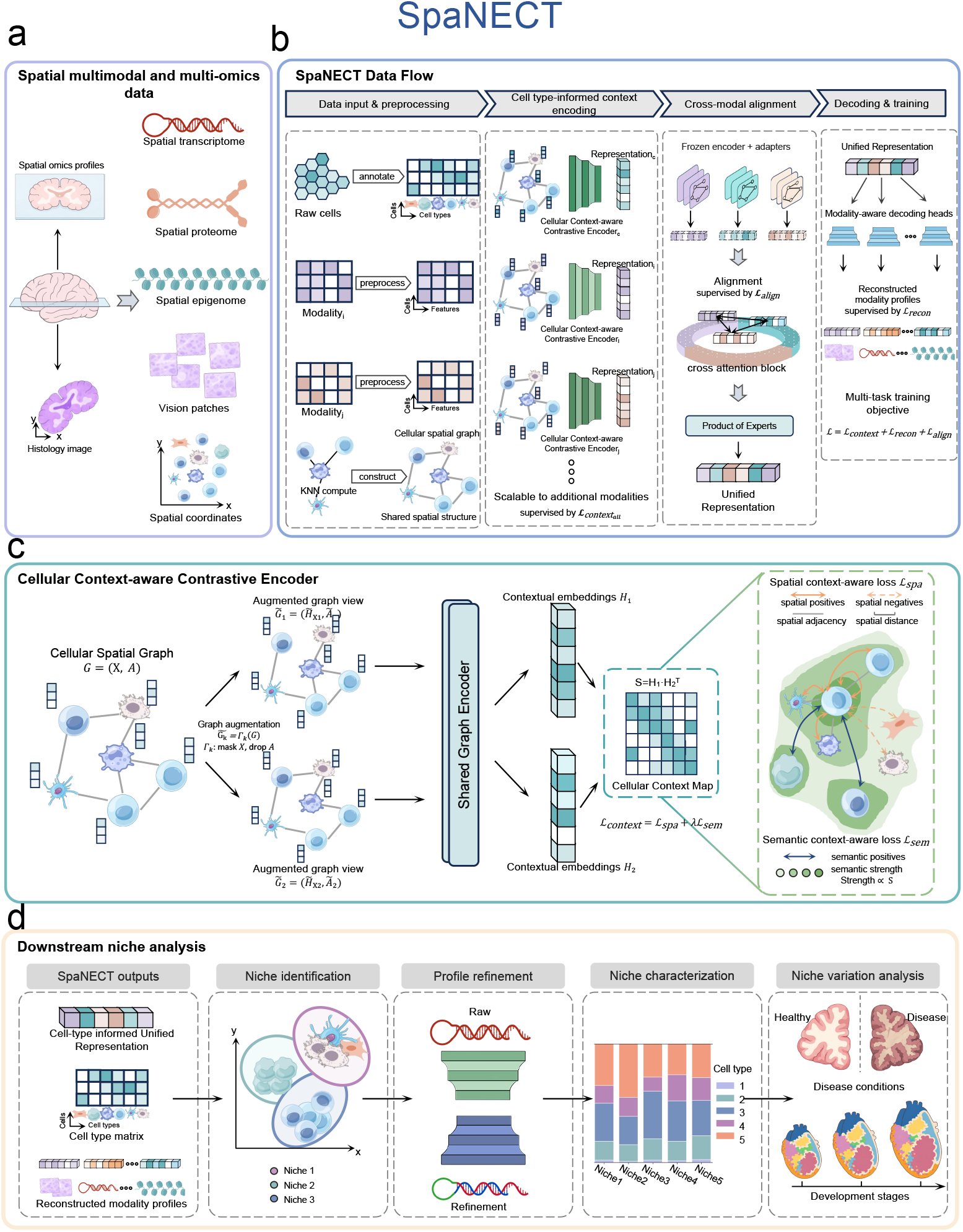
Overview of SpaNECT. (A) Spatial multimodal and multi-omics inputs, including spatial transcriptomic data, additional molecular modalities, histology-derived images, and spatial coordinates. (B) SpaNECT data flow for cell-type-informed multimodal representation learning. Modality-specific feature matrices, reference-informed cell-type information, and a shared cellular spatial graph are used to learn contextual embeddings, which are subsequently aligned and aggregated into a cell-type-informed unified representation. (C) Cellular Context-aware Contrastive Encoder (CCCE). CCCE learns contextual embeddings from augmented graph views while preserving spatial and semantic context consistency. (D) Downstream niche analysis using SpaNECT outputs.

### 2.2 SpaNECT identifies and interprets laminar niches in the human dorsolateral prefrontal cortex from spatial multimodal data

We first evaluated SpaNECT on the canonical 10x Visium human dorsolateral prefrontal cortex dataset[40], which comprises 12 sections from three donors with manual annotations for seven anatomical regions, namely cortical layers 1–6 and white matter (WM). The reference-informed cell-type matrix used by SpaNECT was inferred from an external human brain single-nucleus RNA-seq reference [44]. Using these manual layer annotations[40] as the ground truth, we assessed the agreement between inferred partitions and reference labels with the adjusted Rand index (ARI) and normalized mutual information (NMI), and compared SpaNECT with eight representative spatial clustering and partitioning methods, including DeepST[30], GraphST[25], SEDR[21], STAGATE[20], STAIG[31], stDCL[26], stLearn[28], and CytoCommunity[27]. Across all 12 sections, SpaNECT achieved the highest mean ARI (0.5944) and the highest mean NMI (0.69) (Fig. 2a). For a closer inspection of the inferred spatial partitions, we further focused on two representative sections, #151507 and #151672 (Fig. 2b,c), whereas the corresponding results for the remaining sections are provided in Supplementary Fig. S1. In both sections, SpaNECT recovered the annotated laminar arrangement with high fidelity (ARI = 0.5912 and 0.8353, respectively), and the inferred groups remained well separated in the learned embedding space. On section #151672, SpaNECT further outperformed strong baselines such as DeepST (ARI = 0.7637), stDCL (ARI = 0.6628), and GraphST (ARI = 0.6251). These results indicate that SpaNECT more clearly resolves laminar boundaries, whereas competing methods show layer mixing or positional discrepancies between adjacent layers. To systematically evaluate the contribution of SpaNECT components and auxiliary inputs, we performed ablation studies on the human dorsolateral prefrontal cortex dataset (Supplementary Fig. S2a–c and Supplementary Note 4). The complete model consistently achieved the best overall performance across the 12 tissue sections. Removing image or cell-type inputs, removing key model components, omitting the two-stage training strategy, or replacing the default SPOTlight-derived cell-type matrix led to reduced or less stable performance, supporting the importance of complementary multimodal inputs, modality-specific cellular context learning, cross-modal alignment, and high-quality cell-type information.

**Fig. 2.**
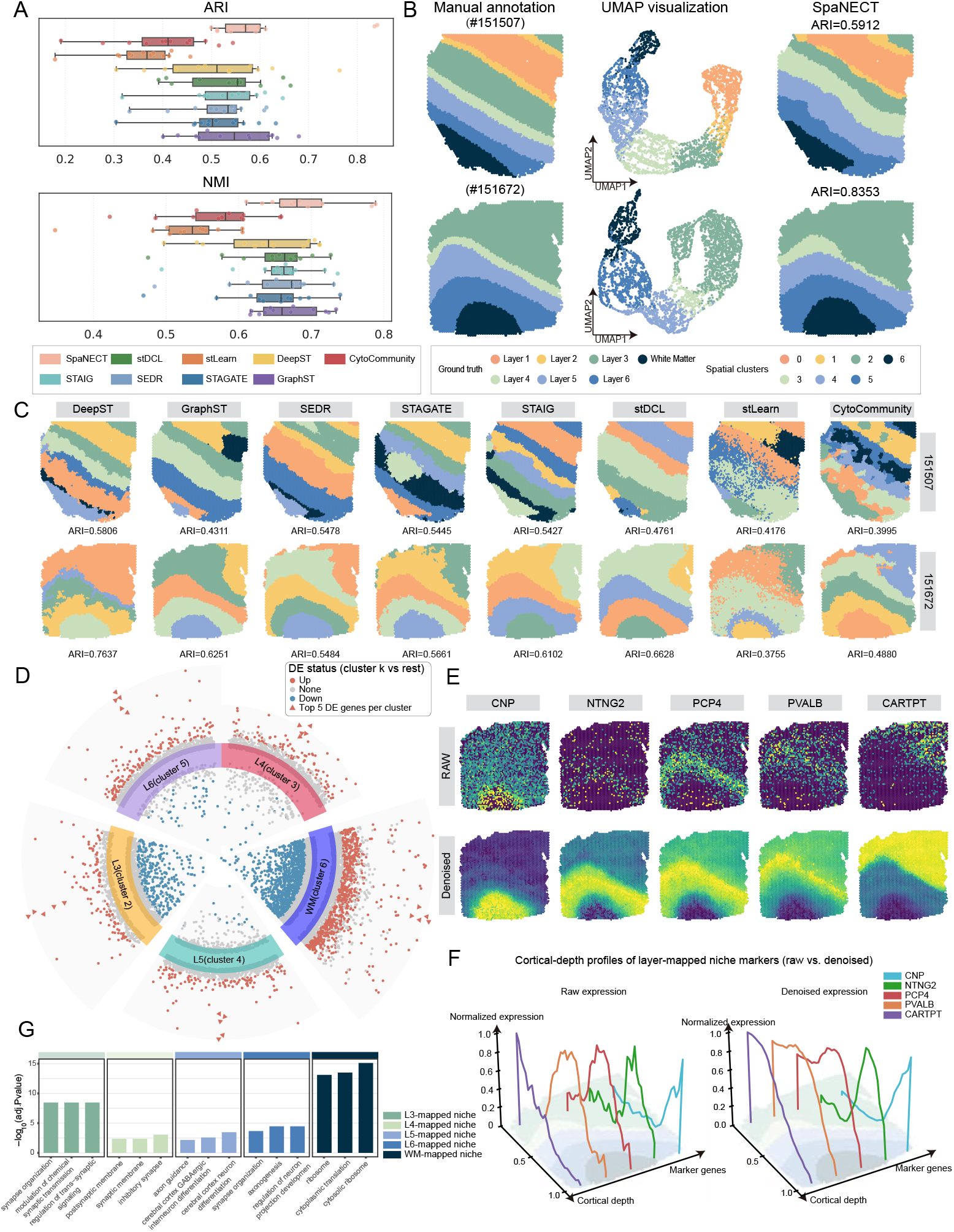
SpaNECT identifies and interprets laminar niches in the human dorsolateral prefrontal cortex from spatial multimodal data. (A) ARI and NMI comparison of SpaNECT and eight baseline methods across all 12 human dorsolateral prefrontal cortex slices. (B) Manual cortical layer annotations, SpaNECT embeddings, and SpaNECT-inferred niches on two representative slices (#151507 and #151672). (C) Spatial clustering results of baseline methods on the same two slices, with ARI values shown. (D) Circular volcano plot of cluster-specific differential expression on slice #151672, highlighting significant genes and top upregulated marker genes. (E) Spatial maps of representative niche-marker genes (*CNP, NTNG2, PCP4, PVALB*, and *CARTPT*) before and after denoising. (F) Cortical-depth expression profiles of the same marker genes before and after denoising across annotated cortical layers. (G) Gene Ontology enrichment analysis of marker genes for layer-mapped niches.

We then asked whether these SpaNECT-inferred partitions could be interpreted as biologically meaningful laminar niches rather than accurate spatial segments alone. To this end, we performed cluster-specific differential expression (DE) analysis on section #151672 by comparing each SpaNECT-inferred cluster against all remaining cells (cluster *k* versus rest; Fig. 2d). The top upregulated marker genes identified for each cluster enabled annotation of the major layer-mapped niches, including L3, L4, L5, L6, and WM. We further visualized representative niche-marker genes, namely *CNP, NTNG2, PCP4, PVALB*, and *CARTPT* (Fig. 2e). Among these markers, *PCP4, NTNG2*, and *CARTPT* corresponded to layer-enriched genes reported in the original study[40], supporting the relevance of the inferred niches to established cortical lamination. *PVALB* is a canonical inhibitory interneuron marker supported by large-scale human cortical cell-type atlases [45], whereas *CNP*, a myelin- and oligodendrocyte-associated gene enriched in white matter [46, 47], was concentrated in the WM-mapped niche. Notably, compared with raw expression, SpaNECT-reconstructed denoised maps displayed clearer layer-restricted spatial distributions, especially for *CNP* in WM and for *NTNG2, PCP4, PVALB*, and *CARTPT* across cortical niches (Fig. 2e). Plotting normalized marker expression along cortical depth provided an orthogonal view of the same trend: denoised signals were more consistent within layers and displayed clearer inter-layer transitions, improving the correspondence between layer-specific peaks and cortical-depth intervals (Fig. 2f).

Finally, Gene Ontology enrichment analysis of niche marker genes further characterized functional differences among the layer-mapped niches, with the top terms shown in Fig. 2g. Specifically, the L3- and L4-mapped niches were enriched for synapse organization, trans-synaptic signaling, synaptic membrane, and inhibitory synapse-related terms; the L5- and L6-mapped niches showed enrichment for axon guidance, axonogenesis, and cortical neuron differentiation-related programs; and the WM-mapped niche was dominated by ribosome, cytoplasmic translation, and cytosolic ribosome signatures. These results further support that SpaNECT-inferred niches capture biologically coherent molecular programs rather than purely geometric partitions.

### 2.3 SpaNECT reveals condition-associated niche reorganization in Alzheimer’s disease from spatial multimodal data

To assess whether SpaNECT could support comparison of niche organization across pathological conditions, we applied it to a disease–control setting in the human dorso- lateral prefrontal cortex, using a control section (CT, 2-5) and an Alzheimer’s disease section (AD, T4857) from age-matched donors [41]. The reference-informed cell-type matrix was constructed using the same external human brain single-nucleus RNA-seq reference as in the preceding analysis [44]. Expert-curated cortical layer annotations spanning Layers 1–6 and white matter (WM) provided a common anatomical reference for cross-condition evaluation (Fig. 3a). Across both sections, SpaNECT showed consistently strong performance relative to representative baselines by ARI, NMI, adjusted mutual information (AMI), and the aggregated overall score (Fig. 3b). It also recovered laminar niches that aligned closely with the annotated cortical architecture, whereas several baseline methods produced more fragmented assignments, particularly near layer boundaries and within the WM region (Fig. 3c).

**Fig. 3.**
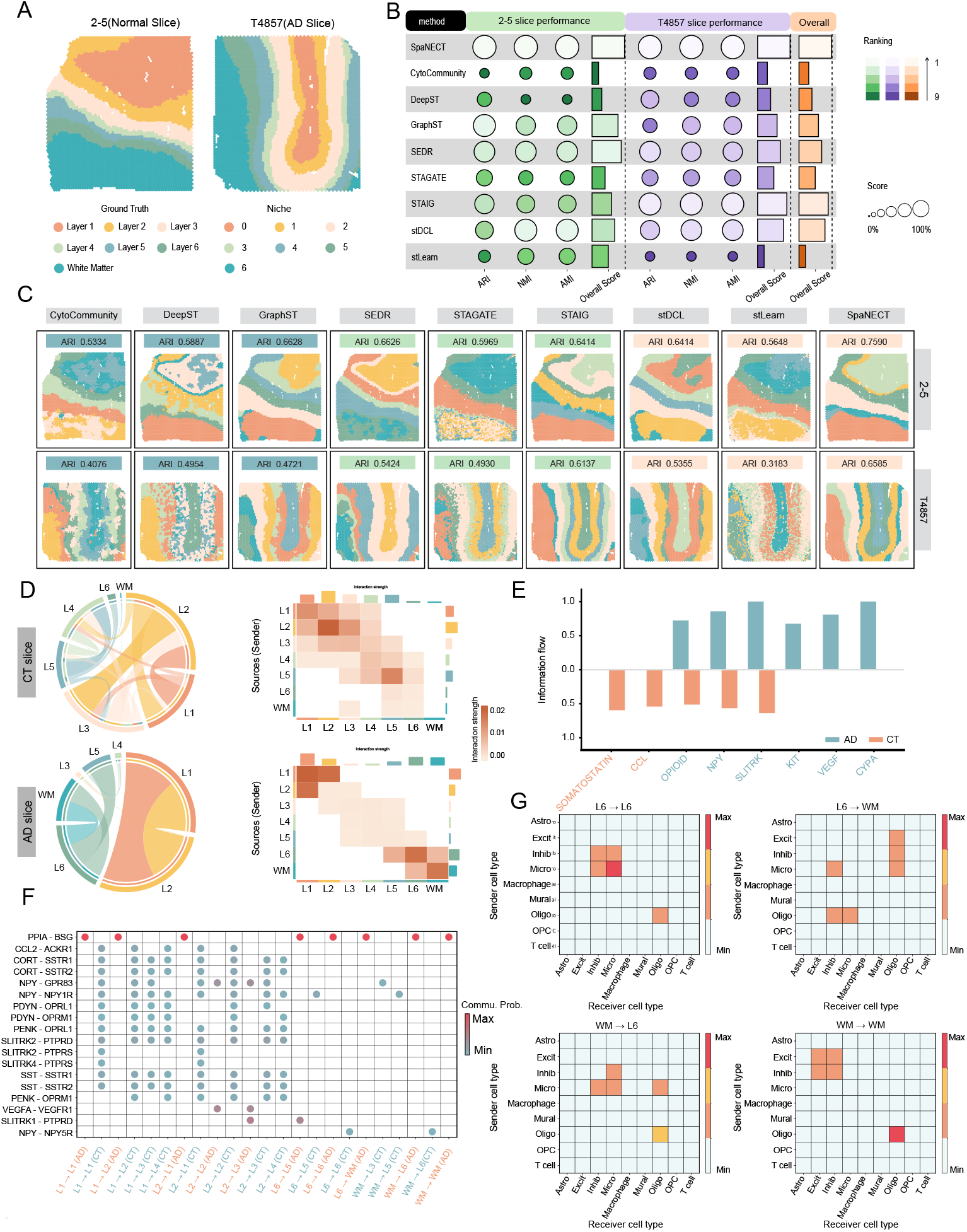
SpaNECT reveals condition-associated niche reorganization in Alzheimer’s disease from spatial multimodal data. (A) Ground-truth cortical layer annotations for the control (CT) and Alzheimer’s disease (AD) sections. (B) Benchmarking performance of SpaNECT and baseline methods on the CT and AD sections, evaluated by ARI, NMI, AMI, and an aggregated overall score. (C) Layer-mapped niche assignments inferred by SpaNECT and baseline methods on the CT section (top) and the AD section (bottom), with the ARI of each method shown in the corresponding panel. (D) Heatmaps and chord diagrams of interaction weights between layer-mapped niches in CT (top) and AD (bottom), showing condition-associated changes in niche-level communication, particularly involving L6 and WM. (E) Pathway-level information flow in CT and AD for representative signaling pathways. (F) Representative condition-associated ligand–receptor pairs contributing to communication between layer-mapped niches. Dot size denotes communication probability, and dot color denotes relative enrichment between CT and AD. (G) Source–target cell-type decomposition of AD-enriched *PPIA*–*BSG* signaling between L6 and WM, shown separately for L6–L6, L6–WM, WM–L6, and WM–WM interactions.

We next compared communication patterns between layer-mapped niches in CT and AD. Comparison of the interaction-weight heatmaps and chord diagrams revealed that interaction strengths in AD were more concentrated on pairs involving deep cortical layers and white matter, with the most prominent increase observed in communication between Layer 6 (L6) and WM (Fig. 3d). This disease-associated increase in communication between L6 and WM suggests that interactions between deep cortical and white-matter compartments may be particularly relevant to AD-associated pathology. Previous studies have likewise reported abnormalities in deep cortical layers and WM in AD, and linked these alterations to disease progression [48, 49].

To further probe the pathological basis of this inter-layer communication shift, we examined pathway-level information flow. Among the representative pathways shown in Fig. 3e, OPIOID, NPY, SLITRK, KIT, VEGF, and CypA displayed higher information flow in AD, whereas SOMATOSTATIN and CCL2 were relatively stronger in CT. Several of these pathways have previously been linked to biological processes relevant to AD, including somatostatin dysregulation, neuropeptide Y-associated synaptic resilience, opioid signaling-related regulation of microglial homeostasis, VEGF-mediated neurovascular responses, and broader neuroinflammatory mechanisms [50–54]. These pathway-level differences suggest that the altered inter-layer communication in AD may involve coordinated changes across multiple disease-relevant signaling programs rather than an isolated perturbation. We then examined condition-associated ligand–receptor pairs mediating communication between layer-mapped niches in the CT and AD sections. We observed several pairs with differential prominence between the two conditions, including *VEGFA*– *VEGFR1, CCL2* –*ACKR1*, and neuropeptide-related interactions such as *SST* –*SSTR, NPY* –*NPYR*, and *PDYN* /*PENK* –*OPRL1* ; among these, *PPIA*–*BSG* signaling was especially prominent in AD (Fig. 3f). The AD-enriched *PPIA*–*BSG* signal was detected across multiple layer-mapped niche interactions and was most pronounced in communication between L6 and WM, suggesting a potential role for this ligand–receptor pair in disease-associated communication reorganization. Previous studies have also implicated CypA (*PPIA*)–CD147 (*BSG*) signaling in inflammatory, neurovascular, and blood–brain barrier-related dysfunction in AD, supporting the potential relevance of *PPIA*–*BSG* to pathological communication changes in the AD cortex [55–57].

Finally, we decomposed the AD-enriched *PPIA*–*BSG* signaling between L6 and WM into source and target cell-type contributions (Fig. 3g). Notably, the contribution pattern was heterogeneous across cell-type pairs. Within the L6-mapped niche, the strongest signal arose from microglia-to-microglia communication, accompanied by additional interactions between microglia and inhibitory neurons, as well as a smaller oligodendrocyte-associated component. Communication from L6 to WM involved multiple distinct components, including excitatory- or inhibitory-neuron-to-oligodendrocyte signaling, as well as interactions from microglia to inhibitory neurons or oligodendrocytes and from oligodendrocytes to microglia or inhibitory neurons. In the reciprocal communication from WM to L6, oligodendrocyte-to-oligodendrocyte signaling was the most prominent component, together with additional contributions from microglia to inhibitory neurons, microglia, or oligodendrocyte-lineage cells. Within the WM-mapped niche, the signal was again dominated by oligodendrocyte-associated self-interactions, accompanied by smaller excitatory- and inhibitory-neuronal components. Overall, this cell-type-resolved decomposition indicates that the AD-enriched *PPIA*–*BSG* signal is concentrated in interactions involving microglia, inhibitory neurons, and oligodendrocyte-lineage cells, with oligodendrocyte-associated components remaining particularly prominent between L6 and WM and within WM. This pattern is consistent with previous reports linking microglia-oligodendrocyte or myelin-associated alterations to AD pathology [49, 58]. Details of definitions, normalization, selection criteria, layer mapping, and the cell-type fraction decomposition are provided in Supplementary Note 7.

### 2.4 SpaNECT delineates spatial niches and regulatory programs in the E15.5 mouse brain from spatial epigenome–transcriptome data

We applied SpaNECT to an E15.5 mouse brain spatial multi-omics dataset with paired spatial gene expression (RNA) and chromatin accessibility (ATAC) profiles(Fig. 4a) [14]. The reference-informed cell-type matrix was derived using a developing mouse brain single-cell RNA-seq reference [59]. The expert anatomical annotation in Fig. 4a provides regional context for interpretation. Under the full configuration (RNA + ATAC + cell type), SpaNECT identified spatially coherent, regionally organized niches with well-defined boundaries (Fig. 4b). Relative to the RNA-only and ATAC-only settings, joint modeling of RNA and ATAC produced more continuous spatial structures, whereas inclusion of the cell-type modality further sharpened niche boundaries in a manner consistent with the major anatomical organization of the section (Fig. 4b). These visual patterns were supported by Moran’s I comparisons across representative baselines, with SpaNECT achieving spatial autocorrelation comparable to that of the strongest unimodal methods while outperforming the compared multimodal methods (Fig. 4c). We then examined the modality weights learned during cross-modal alignment, which reflect the contribution of each modality to the aligned niche representation. The weights varied across niches (Fig. 4d), indicating differential contributions of RNA and ATAC across developing brain regions rather than uniform contributions to niche delineation. RNA markers and niche-enriched spatially variable (SV) peaks further provided interpretable molecular signatures for the inferred niches (Fig. 4e,f). Representative RNA markers highlighted distinct developmental programs, including *Lef1* -associated forebrain and medial pallial signatures together with *Shh*related ventral patterning signals, supporting the biological plausibility of the inferred niche organization [60, 61].

**Fig. 4.**
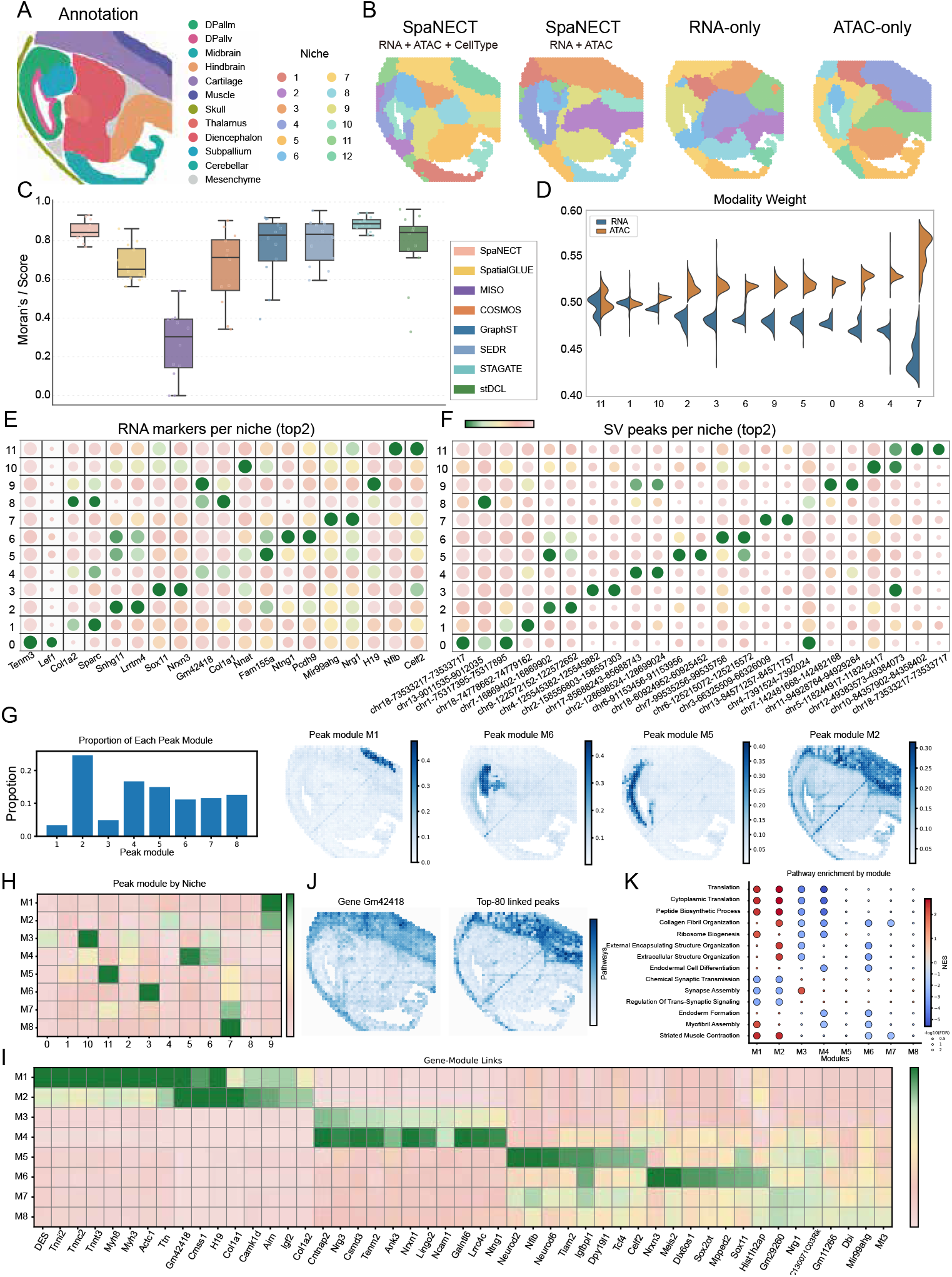
SpaNECT delineates spatial niches and regulatory programs in the E15.5 mouse brain from spatial epigenome–transcriptome data. (A) Reference anatomical annotation of the E15.5 mouse brain section. (B) Spatial niches identified by SpaNECT under the full model (RNA + ATAC + cell type) and modality ablations (RNA + ATAC, RNA-only, and ATAC-only), illustrating the benefit of integrating RNA, ATAC, and cell-type information for spatially coherent niche delineation. (C) Box plots of Moran’s *I* scores for SpaNECT and baseline methods, showing the spatial autocorrelation of inferred niches across spatial single-omics and spatial multi-omics methods. (D) Modality weights of RNA and ATAC, denoting their relative contributions to the aligned niche representation of SpaNECT. (E) RNA marker dot plot across SpaNECT niches. Dot color denotes within-niche mean expression, and dot size denotes the fraction of cells with detectable expression. (F) Spatially variable (SV) peak dot plot across SpaNECT niches. Dot color denotes within-niche mean accessibility, and dot size denotes the fraction of cells with detectable accessibility. (G) Peak-module discovery on selected spatially variable (SV) peaks, showing the proportion of each peak module (left) and spatial activity maps of representative peak modules (right), demonstrating distinct accessibility programs. (H) Heatmap of peak-module enrichment across SpaNECT niches, showing module–niche associations. (I) Gene–module link heatmap derived from gene–peak interaction analysis, summarizing gene–module associations for regulatory interpretation. (J) Representative gene–peak example showing the spatial expression of *Gm42418* (left; z-score) and the aggregated accessibility of its top 80 linked peaks (right; z-score), showing spatial correspondence with the M2-associated region. (K) Module-level pre-ranked pathway enrichment (GSEA) using gene–module association scores as ranking statistics, visualized as a dot plot (color = NES, size = − log_10_ FDR), highlighting distinct functional themes across peak modules.

To characterize these niches from a regulatory perspective, we performed peak-module discovery on selected spatially variable (SV) peaks and identified modules of varying sizes, each exhibiting distinct region-specific accessibility patterns (Fig. 4g). Mapping these modules back to SpaNECT niches revealed clear module-niche associations, indicating that different niches are associated with distinct accessibility programs (Fig. 4h). We then used gene-peak interaction analysis to summarize module-associated genes and thereby facilitate interpretation of these accessibility-defined modules (Fig. 4i). Notably, genes strongly associated with specific modules included developmental regulators such as *Lef1* and *Shh*, consistent with their established roles in forebrain patterning and regionalization [60, 61]. To visualize this regulatory correspondence more directly in tissue space, we selected *Gm42418* from M2 along with its top 80 linked peaks and compared their spatial patterns. The spatial distribution of *Gm42418* closely matched the aggregated accessibility of its linked peaks and coincided with the M2-associated region (Fig. 4j). Module-level pre-ranked pathway enrichment analysis further resolved distinct functional themes across modules (Fig. 4k) [62]. Translation-related programs, including translation, cytoplasmic translation, peptide biosynthetic process, and ribosome biogenesis, were enriched in both M1 and M2, whereas synaptic and neuronal signaling programs, including chemical synaptic transmission, synapse assembly, and regulation of trans-synaptic signaling, were most prominent in M3. Extracellular matrix and structural organization terms, such as collagen fibril organization, external encapsulating structure organization, and extra-cellular structure organization, were most prominent in M2. Muscle-related programs, including myofibril assembly and striated muscle contraction, were also enriched in both M1 and M2. These analyses show that SpaNECT resolves niche-associated accessibility modules and links them to spatially localized regulatory programs in the developing brain. Computational details of SV peak selection, peak-module construction, gene–peak scoring, and module-level enrichment analysis are provided in Supplementary Note 8.

### 2.5 SpaNECT resolves and characterizes heterogeneous spatial niches in the human tonsil from spatial RNA–ADT data

We next applied SpaNECT to a human tonsil spatial multi-omics dataset with paired RNA and antibody-derived tag (ADT) data (Fig. 5a) [42, 63]. The reference-informed cell-type matrix was obtained using a human tonsil single-cell atlas as the reference [64]. This tissue section exhibits a highly heterogeneous lymphoid architecture, where B-cell follicles are embedded in T-cell-enriched regions and further bounded by epithelial and connective tissues. Manual annotation defined four major anatomical regions, namely connective & epithelial tissue, germinal center (GC), lymphoid follicle, and tonsillar parenchyma, which served as a reference for evaluation (Fig. 5a). Under the full model (RNA + ADT + cell type), SpaNECT identified four spatially coherent niches that matched these major structures well (Fig. 5b). The full configuration out-performed the RNA + ADT setting and both unimodal ablations, and SpaNECT also achieved the highest ARI among the compared single-omics and multi-omics methods (Fig. 5b,c), indicating that paired transcriptomic and protein profiles, together with cell-type information, provide a stronger basis for robust niche delineation in this structurally complex immune tissue [65–67].

**Fig. 5.**
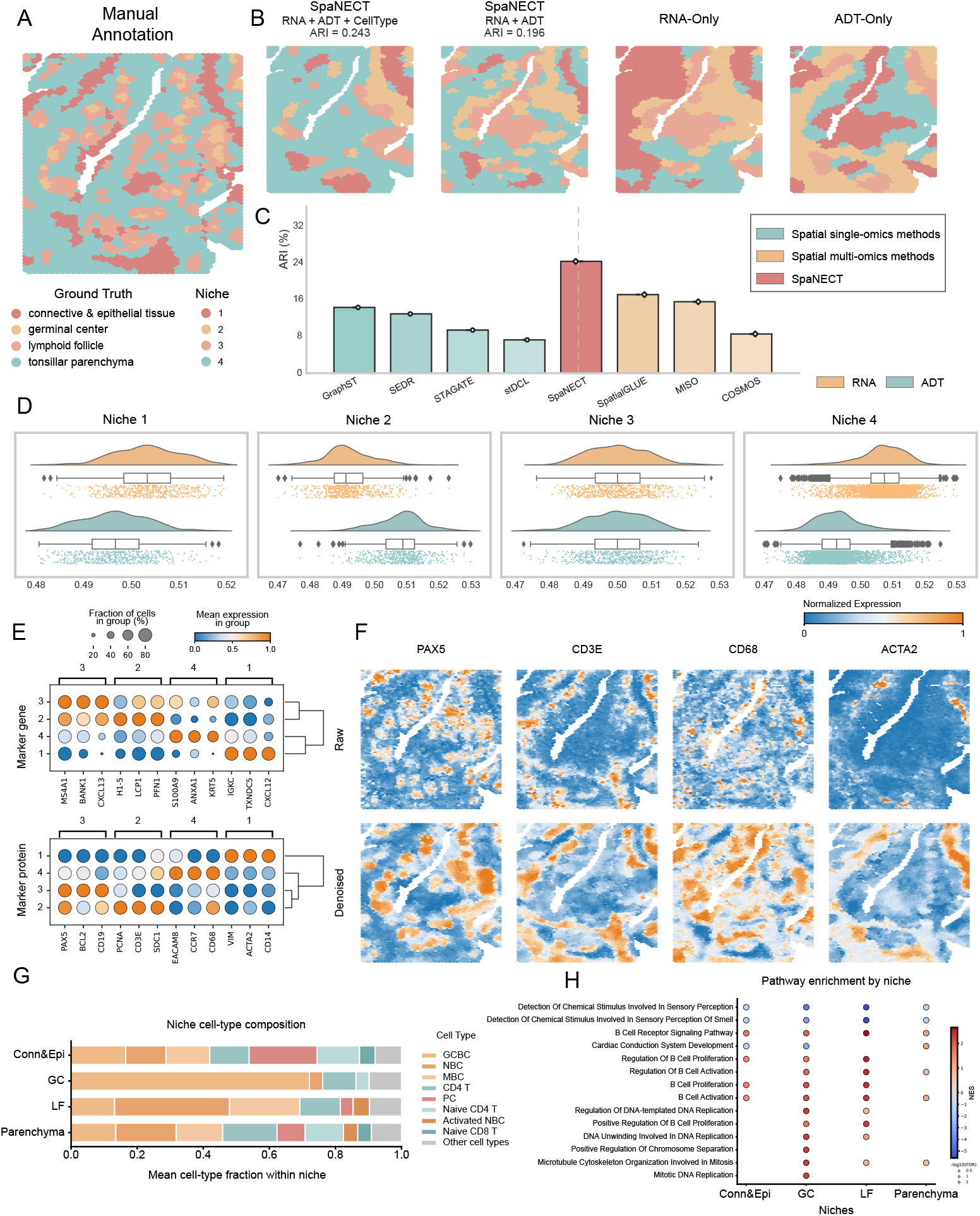
SpaNECT resolves and characterizes heterogeneous spatial niches in the human tonsil from spatial RNA–ADT data. (A) Manual anatomical annotation of a human tonsil section, defining four major anatomical regions: connective & epithelial tissue, germinal center, lymphoid follicle, and tonsillar parenchyma. (B) Spatial niches identified by SpaNECT under the full model (RNA + ADT + cell type) and modality ablations (RNA + ADT, RNA-only, and ADT-only), with niche identification accuracy quantified by adjusted Rand index (ARI). (C) Quantitative comparison of niche identification accuracy (ARI) between SpaNECT and representative competing methods from spatial single-omics and spatial multi-omics analysis. (D) Modality weights learned during cross-modal alignment, shown as the per-spot distributions of RNA and ADT contributions within each SpaNECT niche; higher weights indicate greater contribution of the corresponding modality to the aligned niche representation. (E) Representative RNA and ADT markers across niches. Dot plots summarize within-niche mean expression (color) and the fraction of spots expressing each marker (dot size). (F) Raw and denoised spatial maps of representative ADT markers. (G) Cell-type composition of each niche shown as stacked bar plots. (H) Niche-specific pathway enrichment analysis based on ranked differential expression from RNA data (each niche versus all other spots). Dot color denotes normalized enrichment score (NES), and dot size denotes significance (− log_10_ FDR).

We then examined the modality weights learned during cross-modal alignment, which reflect the contribution of each modality to the aligned niche representation (Fig. 5d). The weights varied across niches, indicating differential contributions of RNA and ADT rather than uniform contributions across the tissue. Specifically, RNA contributed more strongly to the connective & epithelial niche and the tonsillar parenchyma niche, ADT showed a stronger contribution in the germinal-center niche, whereas the lymphoid-follicle niche exhibited a more balanced contribution profile (Fig. 5d). Representative RNA and ADT markers further supported these niche assignments (Fig. 5e). B-lineage RNA markers, including *PAX5, BCL2, MS4A1, BANK1, IGKC*, and *CXCL13*, together with the ADT marker CD19, were concentrated in the germinal-center and lymphoid-follicle niches, whereas *CD3E* and *CCR7* localized more strongly to the parenchyma-like niche, and myeloid- or stromal-associated markers such as CD68 and *ACTA2* highlighted the connective & epithelial niche. Given the inherently low signal-to-noise characteristics of spatial omics, particularly in modalities beyond transcriptomics [34, 68], we further compared representative protein maps before and after denoising. SpaNECT denoising enhanced spatial coherence and made these niche-resolved ADT patterns more distinct than the raw signal while preserving the major tissue layout (Fig. 5f) [69–72].

These niches were further distinguished by their cellular composition and functional programs. The germinal-center niche was dominated by GC-associated B cells, the lymphoid-follicle niche showed broader B-cell enrichment, the parenchyma niche contained larger T-cell fractions, and the connective & epithelial niche exhibited a more mixed composition (Fig. 5g). At the functional level, pathway enrichment based on ranked differential expression further separated these niches into biologically consistent programs (Fig. 5h). The germinal-center and lymphoid-follicle niches were enriched for B-cell activation, B-cell receptor signaling, and proliferation-related programs, with the germinal-center niche additionally showing DNA replication and mitotic terms consistent with active germinal-center reactions [73–75]. By contrast, the non-lymphoid niches displayed distinct interface- and immune-associated functional themes consistent with their spatial positions and mixed immune composition (Fig. 5h).

### 2.6 SpaNECT tracks maturation-associated niche reorganization in embryonic chick heart from time-series spatial multimodal data

To investigate how spatial niches are remodeled during chick heart development, SpaNECT was applied to an embryonic chick heart developmental series profiled at Day 7 (D7), Day 10 (D10), and Day 14 (D14), encompassing a period of pronounced chamber maturation and structural reorganization [43]. For this dataset, the cell-type matrix used by SpaNECT was constructed from the cell-type annotations provided in the original developing chick heart study. The anatomical annotations in Fig. 6a provide a developmental reference for these sections. SpaNECT identified spatially coherent niches that closely recapitulated the major cardiac compartments at each stage (Fig. 6b). Across all three stages, SpaNECT achieved the highest agreement with the anatomical annotations among the compared methods, as quantified by ARI (Fig. 6c), indicating robust niche identification despite substantial developmental variation. Representative region-to-niche correspondences further showed that key structures, including the atria, valves, and trabecular LV/endocardium-associated region, remained readily traceable in the SpaNECT niche maps across D7–D14 (Fig. 6d), enabling cross-stage comparison of niche organization.

**Fig. 6.**
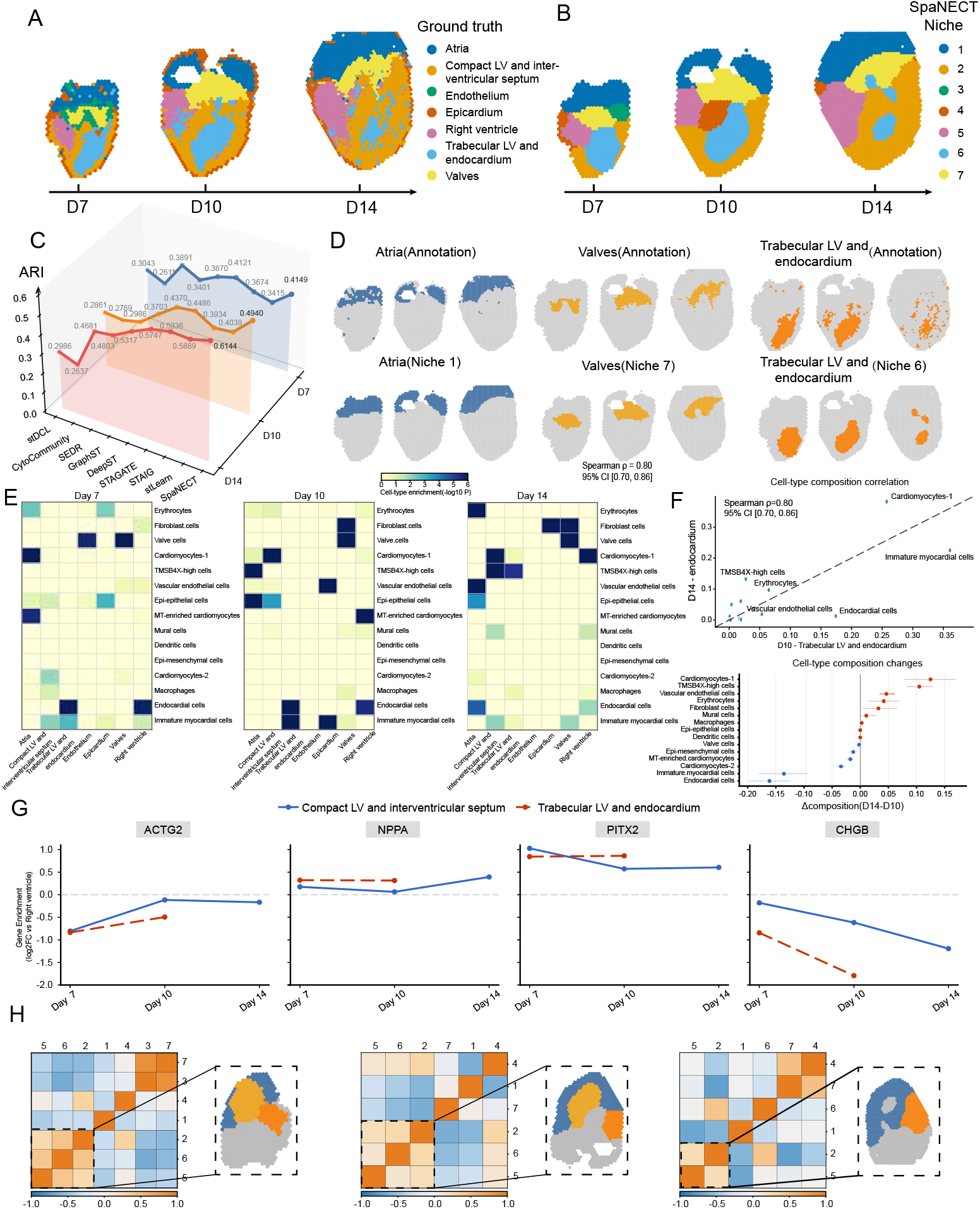
SpaNECT tracks maturation-associated niche reorganization in embryonic chick heart from spatial multimodal data. (A) Anatomical annotations for chick heart sections profiled at D7, D10, and D14. (B) Spatial niches identified by SpaNECT in the three sections. (C) Quantitative comparison of niche identification performance across developmental stages, measured by adjusted Rand index (ARI), for SpaNECT and competing methods. (D) Representative examples showing the correspondence between selected annotated regions and their matched SpaNECT niches across D7–D14 (highlighted region in color; remaining spots in gray). (E) Cell-type enrichment landscapes across SpaNECT niches and developmental stages, shown as niche-by-cell-type enrichment heatmaps for D7, D10, and D14. (F) Comparison between the SpaNECT niche mapped to the D10 trabecular LV/endocardium region and the D14 endocardium-associated niche, showing correlation of cell-type profiles (top) and changes in cell-type composition (bottom). (G) Marker-gene enrichment trends for SpaNECT niches mapped to the compact LV and interventricular septum and trabecular LV/endocardium across development, computed relative to the SpaNECT niche mapped to the right ventricle (RV). Because the trabecular LV/endocardium-associated niche is no longer detected at D14, enrichment trends for this niche are shown only for D7 and D10. (H) Cross-stage niche similarity analysis, shown as correlation heatmaps highlighting highly correlated SpaNECT niches across developmental stages together with their spatial localization.

We next characterized SpaNECT niches from a cell-type composition perspective across developmental stages (Fig. 6e). The resulting enrichment patterns delineated stage-resolved niche identities consistent with cardiac compartment biology. Atria-associated niches were dominated by cardiomyocyte populations, including MT-enriched cardiomyocytes, whereas niches mapped to the compact LV and interventricular septum were enriched for cardiomyocytes together with immature myocardial cells, consistent with ongoing ventricular maturation. Endothelium-associated niches showed enrichment of vascular endothelial cells together with mural cells, and valve-associated niches displayed selective enrichment of valve cells with stromal components such as fibroblast cells, in line with cushion remodeling and endothelial-to-mesenchymal transition during valve morphogenesis [76–79]. These niche-specific cell-type profiles suggest that chick heart maturation is accompanied by coordinated reorganization of compartment-associated cellular assemblies, particularly in ventricular- and valve-associated regions. To interrogate late-stage remodeling of trabecular-associated structures, we compared the niche mapped to the D10 trabecular LV/endocardium region with the D14 endocardium-associated niche (Fig. 6f). Their cell-type profiles were strongly concordant (Spearman *ρ* = 0.80; bootstrap 95% CI [0.70, 0.86]; Fig. 6f), indicating that the overall cell-type composition of these niches remains similar across stages and reflects a continuous developmental progression. Examining the change in cell-type composition (Δ = D14 − D10) further revealed more specific shifts: Cardiomyocytes-1, *TMSB4X* -high cells, and vascular endothelial cells increased, whereas immature myocardial cells and endocardial cells decreased (Fig. 6f). These findings are consistent with current understanding that endocardial–myocardial signaling, including Notch and neuregulin pathways, coordinates trabecular remodeling during maturation and is accompanied by progressive refinement of myocardial states and endothelial programs; the cell types highlighted above may therefore contribute to this remodeling process [80].

To provide complementary molecular support for ventricular-associated niches, we examined relative expression trends of ACTG2, NPPA, PITX2, and CHGB, using the niche mapped to the right ventricle as the reference (Fig. 6g). We observed coherent, niche-dependent temporal patterns, including persistent right-ventricle-associated enrichment of *CHGB* and left-sided enrichment of *PITX2*, consistent with established chamber-associated and left–right patterning programs during heart development [43, 81–83]. We then assessed cross-stage niche comparability using correlation-based niche similarity matrices and spatial back-mapping of highlighted niche pairs (Fig. 6h). This analysis identified stable, highly correlated niche correspondences across stages, while also revealing selective reorganization among ventricular-associated niches, consistent with attenuation of trabecular-associated similarity by D14 [43, 79, 80].

## 3 Discussion

SpaNECT is a unified and flexible framework for cell-type-informed spatial niche analysis in spatial multimodal and multi-omics data. SpaNECT separates cellular context learning from multimodal alignment. This design preserves the effectiveness of local cellular context modeling while improving the stability of multimodal alignment across data modalities with distinct noise structures and information content. In addition, the explicit incorporation of cell-type information facilitates biologically grounded niche characterization and comparison and makes the framework more adaptable to increasingly diverse spatial data.

We evaluated SpaNECT across diverse biological contexts, spanning multiple tissues, conditions, developmental stages, and spatial multimodal or multi-omics data types[14, 40–43]. Across these analyses, SpaNECT consistently supported niche identification, multimodal characterization, and comparison of niche organization. At present, SpaNECT remains limited in two respects. Although SpaNECT is designed as a unified and extensible framework for spatial multimodal and multi-omics data, currently available datasets are still constrained by existing profiling technologies and experimental conditions. In many cases, only a limited number of omics layers can be profiled from a given section, whereas complementary modalities are obtained from separate, often adjacent, sections and then computationally aligned for joint analysis[42, 84, 85]. This can limit comprehensive niche characterization. In addition, our experiments indicate that the performance and interpretability of SpaNECT can be influenced, to some extent, by the availability and quality of cell-type annotations.

As spatial profiling technologies continue to generate increasingly diverse data on tissue organization, SpaNECT may support broader applications and provide deeper insights into the organization, function, and variation of cellular niches across tissues and conditions.

## 4 Methods

### 4.1 Data description

We used publicly available spatial multimodal and multi-omics datasets from multiple tissues, disease conditions, and developmental stages to evaluate SpaNECT (Supplementary Table 1). The human dorsolateral prefrontal cortex dataset was generated using the 10x Visium platform and includes 12 sections with 3,460–4,789 spots per section, each with manual cortical layer annotations. For disease-associated niche analysis, we used one control section and one Alzheimer’s disease section from human cortex with corresponding layer annotations. The E15.5 mouse brain dataset was generated using the MISAR-seq platform and contains paired RNA and ATAC profiles from 1,949 spots. The human tonsil dataset was obtained from SpaMosaic and contains paired spatial RNA and ADT profiles from 4,518 spots with manual anatomical annotations. The embryonic chick heart dataset spans three developmental stages, Day 7, Day 10, and Day 14, with 494, 1,039, and 1,967 spots, respectively. Manual or reference annotations were obtained from the corresponding original studies when available.

### 4.2 Data preprocessing

We preprocess each modality to obtain a collection of feature matrices 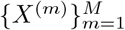, where 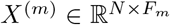 represents the features of modality *m, N* denotes the number of cells, and *F*_*m*_ is the dimensionality of modality-*m* features.

#### RNA (gene expression)

The raw transcript count matrix is preprocessed using Scanpy [86]. Library-size normalization to a target sum of 10^4^ per cell is applied, followed by a log_2_(1+*x*) transformation to stabilize variance. Highly variable genes are then identified using the Seurat v3 strategy [87], and the top 3,000 genes are retained by default, yielding the RNA feature matrix *X*^(rna)^ ∈ ℝ^*N* ×3000^, where *N* denotes the number of cells.

#### Other omics (ATAC/ADT)

For additional molecular modalities available in spatial multi-omics datasets, an analogous normalization-and-transformation procedure is applied to obtain numerically stable feature matrices. The raw ATAC peak matrix or protein/ADT intensity matrix is normalized to a common scale (e.g., total-count normalization) and transformed with log(1 + *x*) when appropriate, producing 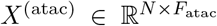 or 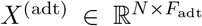. Dataset-specific settings and additional preprocessing details are provided in Supplementary Note 2.

#### Histology image (H&E)

When histological images are available, we crop a local patch from the H&E image centered at each cell and extract a fixed-length visual representation as *X*^(img)^. We adopt BYOL [88] for self-supervised visual representation learning, an advanced consistency-based paradigm that learns discriminative features without manual labels and improves robustness to staining variation and local perturbations; patch size, resolution, data augmentations, and BYOL training details are described in Supplementary Note 5.

### 4.3 Cell-type information inference

Cell-type compositions are estimated with SPOTlight [89]. For each cell, we infer a cell-type proportion matrix *C* ∈ ℝ^*N* ×*K*^, where *N* denotes the number of cells and *K* denotes the number of cell types. The matrix *C* is row-normalized such that each row sums to one. It represents the relative composition of cell types for each cell and serves as the cell-type feature matrix *X*^(ct)^ in SpaNECT.

To assess the effect of the deconvolution choice, additional experiments are conducted using alternative cell-type inference methods, including CARD-free [90] and GraphST [25]. Results from these variants are reported in Supplementary Note 6.

### 4.4 Construction of the cellular spatial graph

For each tissue slice, we encode spatial proximity as an undirected cellular spatial graph *G* = (*V, E*). Let *N* denote the number of cells and let *P* = [*p*_1_, …, *p*_*N*_]^⊤^ ∈ ℝ^*N* ×2^ denote their 2D coordinates, where *p*_*i*_ is the coordinate of cell *i*. We define the node set *V* = 1, …, *N* as the collection of all cells in the tissue slice. The edge set *E* is constructed from spatial neighborhoods computed in the coordinate space. Specifically, for each cell *i*, we identify its *k* nearest neighbors 𝒩_*k*_(*i*) according to Euclidean distances between coordinates in *P*, and form a binary adjacency matrix *A* ∈ {0, 1}^*N* ×*N*^ :

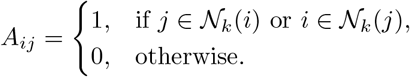

To retain self-information and stabilize message passing, we add self-loops and apply symmetric normalization:

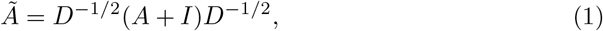

where *D* is the degree matrix of (*A* + *I*) and *I* is the identity matrix. The resulting cellular spatial graph is shared across all modality branches to provide a common topology for learning cellular context representations. In practice, *k*-NN is computed using tree-based neighbor search, and an optional radius-based graph can be used when a distance threshold is preferred.

### 4.5 Cellular Context-aware Contrastive Encoder

#### Encoder architecture

Given a shared cellular spatial graph *G* = (*V, E*) with normalized adjacency *Ã* and modality-specific features 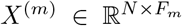, we train an independent Cellular Context-aware Contrastive Encoder (CCCE) branch for each modality *m* ∈ {1, …, *M*} to learn modality-specific cellular context representations. A message-passing graph encoder 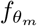 integrates node attributes with spatial adjacency to produce contextual embeddings:

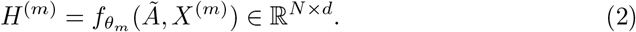

A projection head 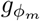 then maps the contextual embeddings into a contrastive space:

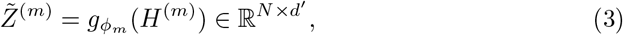

where 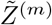 denotes the projected embeddings and *d*^′^ is the projection dimension.

#### Dual-view augmentations

To improve robustness to measurement noise and uncertainty in local spatial neighborhoods, CCCE constructs two randomly augmented views 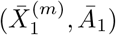 and 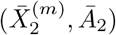 for each modality. Specifically, we corrupt a subset of feature dimensions via feature masking and remove a subset of spatial edges via edge dropping, encouraging the encoder to learn stable cellular context representations.

#### Cellular context map

For the two augmented views, the encoder and projection head produce 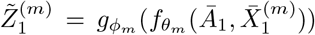 and 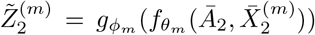. Since CCCE aims to learn a similarity structure that captures cellular context, we apply row-wise *ℓ*_2_ normalization to the projected embeddings to place them on the unit hypersphere, enabling cosine similarity computation by inner products:

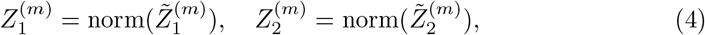

where norm(·) denotes row-wise *ℓ*_2_ normalization. We then define the cellular context map as the cross-view cosine similarity (node similarity) matrix:

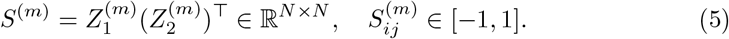

To introduce a differentiable sparsity gate, we further define

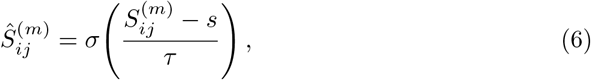

where *s* ∈ [0, 1] is a similarity threshold, *τ >* 0 is the temperature, and *σ*(·) is the sigmoid function.

#### Composite cellular context-aware contrastive objective

CCCE is optimized with a composite context-aware objective consisting of two complementary terms:

i. Spatial context loss 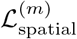. We encourage consistency for the same cell across views and for spatially adjacent cell pairs. The self-alignment term maximizes diagonal similarities:

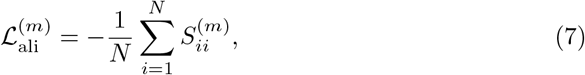

and the neighbor alignment term maximizes similarities over spatial edges:

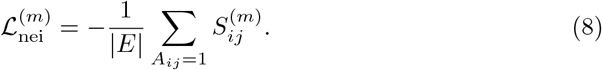 We combine them as:

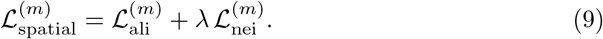
ii. Semantic context loss 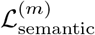. To avoid indiscriminately separating all non-adjacent cell pairs and disrupting latent semantic structure, we impose a semantic sparsification regularizer on disconnected pairs so that only a small subset of semantically plausible non-neighbors can maintain high similarity. Concretely, we apply a sparsity penalty on *Ŝ*^(*m*)^ over pairs with *A*_*ij*_ = 0:

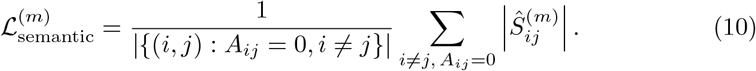 Accordingly, the CCCE context-aware loss for modality *m* is defined as:

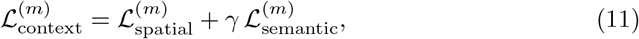

where *λ* and *γ* are hyperparameters controlling the strengths of neighborhood alignment and semantic sparsification, respectively. The overall context-aware loss aggregates across modalities:

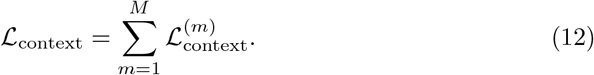

### 4.6 Cross-modal alignment and unified representation

#### Frozen encoders with parameter-efficient adapters

During cross-modal alignment, we reuse the modality-specific graph encoders pretrained in the CCCE stage and freeze their parameters to preserve the unimodal cellular-context structures, thereby minimizing interference from cross-modal optimization. Let the shared spatial graph be *G* = (*V, E*) with normalized adjacency *Ã*. For modality *m*, given node features 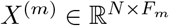, the frozen encoder produces node embeddings

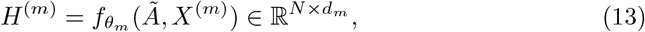

where *N* is the number of cells, *F*_*m*_ is the input feature dimension of modality *m, θ*_*m*_ denotes the fixed parameters of the modality-specific graph encoder, and *d*_*m*_ is the output embedding dimension. To enable parameter-efficient cross-modal alignment while ensuring semantic comparability across modalities, we attach a lightweight adapter 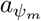 after each frozen encoder to project modality-specific embeddings *H*^(*m*)^ into a shared alignment space of dimension *d*:

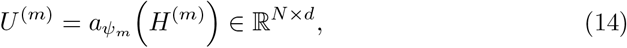

where *ψ*_*m*_ denotes the learnable adapter parameters. Consequently, only *ψ*_*m*_ along with the parameters of subsequent alignment and fusion modules are updated during this stage, while *θ*_*m*_ remains strictly frozen.

#### Cross-attention alignment

Given node embeddings in the shared alignment space 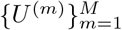, we perform node-wise cross-modal attention to learn soft matching across modality-specific representations. For node *i*, let 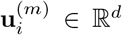 denote its embedding from modality *m*. For each modality *m*, we treat 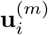 as the query input and use the set 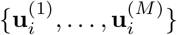 at the same node *i* as candidates for keys and values. This yields a modality-specific attention distribution over other modalities at node *i*, capturing how information from modality *m* attends to representations across all *M* modalities. Specifically, modality-dependent linear projections 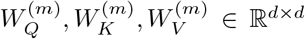 generate queries, keys, and values:

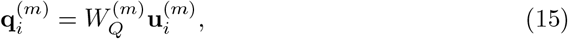

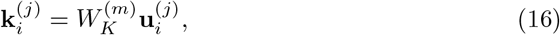

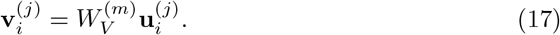

Here *j* indexes the candidate modalities, with *j* = 1, …, *M*. The resulting cross-modal attention weights and aligned node embedding for modality *m* are computed as:

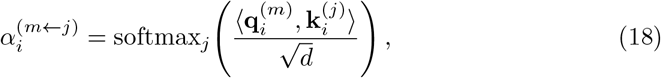

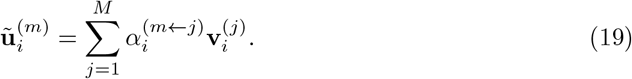

Applying this process for all *m* = 1, …, *M* yields the set of aligned representations 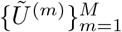, where 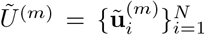. In downstream analysis, the attention weights 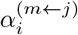 provide an interpretable summary of cross-modal attention and can be aggregated across query modalities to derive node-level modality contribution profiles.

To calibrate the agreement between query representations and their attention-aligned counterparts, we introduce a matched-score alignment regularizer. For each modality *m* and cell *i*, we first define a temperature-scaled matched score:

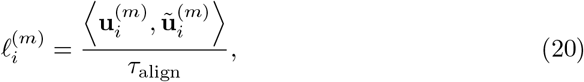

where *τ*_align_ is a temperature parameter. The matched scores are normalized across cells within the same modality:

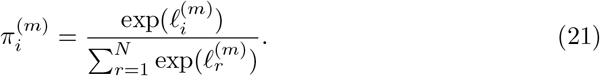

We regularize the resulting matched-score distribution using a uniform target over cells:

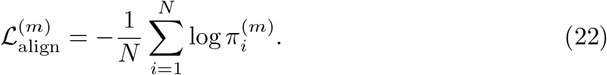

This term provides a lightweight calibration constraint on the matched alignment scores, reducing the tendency of the auxiliary alignment signal to be dominated by a small subset of cells with disproportionately high scores. The final alignment regularization term is averaged across modalities:

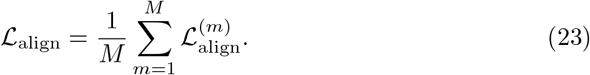

#### Unified representation aggregation

The aligned representations 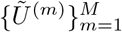 are aggregated into a unified latent representation *Z* ∈ ℝ^*N* ×*d*^ using a Gaussian product-based aggregation scheme. For cell *i*, each modality defines a diagonal Gaussian representation

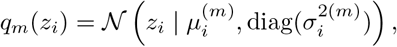

where 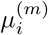 is given by 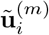, and 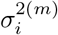 denotes the modality-specific diagonal variance. The aggregated posterior remains Gaussian and has a closed-form precision-weighted solution:

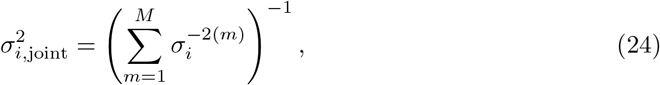

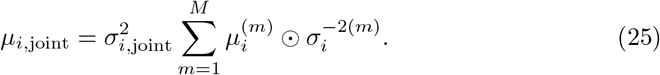

We use *µ*_*i*,joint_ as the unified embedding *z*_*i*_ and retain log 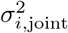 for KL regularization.

#### Modality-aware decoders

Given the unified latent representation *Z* = [*z*_1_, …, *z*_*N*_]^⊤^ ∈ ℝ^*N* ×*d*^, we assign a dedicated decoding head 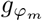 to each modality *m* ∈ {1, …, *M*} to reconstruct modality-specific observations while preserving their distinct structural and statistical characteristics. Each head maps *Z* nonlinearly to the native feature space of modality *m*:

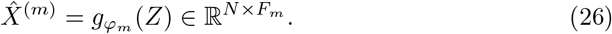

Each 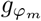 is realized as a two-layer fully connected multilayer perceptron (MLP). Let ℳ_dec_ ⊆ {1, …, *M*}denote the subset of modalities selected for reconstruction supervision; the corresponding reconstruction loss is defined as the mean squared error (MSE) averaged over cells and features:

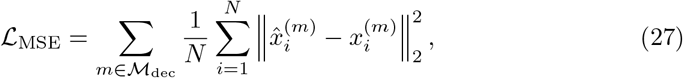

where 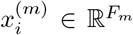 and 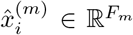 denote the observed and reconstructed feature vectors for cell *i* in modality *m*, respectively. To regularize the unified representation and support a variational interpretation of the aggregated posterior 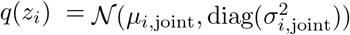, we impose a KL divergence penalty against a standard normal prior:

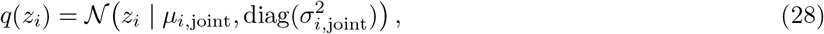

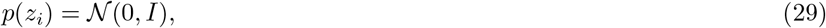

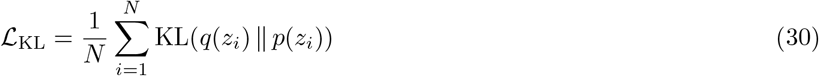

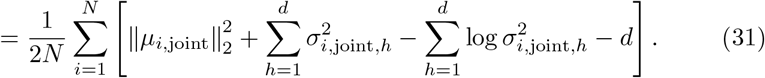

Here, *d* denotes the dimensionality of the latent embedding *z*_*i*_, and *h* indexes latent dimensions. The full reconstruction objective is then formulated as a weighted sum:

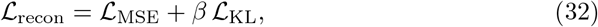

where *β >* 0 is a hyperparameter controlling the strength of latent regularization. Together with the cross-modal alignment loss ℒ_align_ and the context-aware learning term ℒ_context_, ℒ_recon_ forms the overall training objective described in the next section.

### 4.7 Training objective and implementation

We employ a two-stage training strategy: Stage I independently trains the CCCE encoders for each modality to learn stable spatial-context structures, while Stage II freezes the CCCE encoders and optimizes only the lightweight adapters, cross-modal alignment block, latent aggregation, and modality-specific decoders to obtain a unified representation. The overall objective is

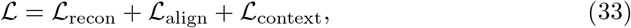

optimized with Adam [91] using an initial learning rate of 5 × 10^−4^ and a weight decay of 1 × 10^−5^. The multimodal stage is trained for 300 epochs; additional implementation details and hyperparameter selection are provided in the Supplementary Information. All experiments were conducted on an NVIDIA RTX 4090 GPU installed in a Linux system with an Intel Xeon Platinum 8370C CPU (2.80 GHz) and 64 GB of memory, and we report software versions and key configurations for reproducibility.

### 4.8 Niche identification by clustering

After model training, the unified representation *Z* was used for niche identification. Initial niche labels were obtained by clustering the learned cell embeddings using K-means [92] by default, with Mclust [93] used as an alternative clustering algorithm when required by the analysis setting. For benchmark datasets with manual annotations, the number of clusters was set to the number of annotated tissue regions or niches. To improve spatial coherence and reduce isolated assignments, we further applied a spatial refinement step based on neighborhood information from the shared cellular spatial graph, and the refined labels were used as the final niche assignments.

### 4.9 Comparative methods and benchmarking setup

We compared SpaNECT with eleven baseline methods spanning two categories: spatial single-omics methods and spatial multi-omics methods. The spatial single-omics baselines included GraphST [25] (v1.1.1), SEDR [21] (v1.0.0), STAGATE [20] (PyG implementation, v1.0.0), stDCL [26] (v1.0.1), stLearn [28] (v1.3.0), DeepST [30] (v2.0.2), CytoCommunity [27] (v1.0.0), and STAIG [31] (v1.0.0). The spatial multi-omics baselines included SpatialGlue [36] (v1.1.5), MISO [33] (v0.1.0), and COSMOS [37] (v1.0). For paired RNA + H&E datasets, we primarily compared SpaNECT against the spatial single-omics baselines, whereas for paired spatial multi-omics datasets, we additionally included the spatial multi-omics baselines for comparison. All competing methods were run using their recommended preprocessing steps and default or author-recommended hyperparameter settings whenever possible. To ensure a fair comparison, the same input data partitions and annotation references were used across methods within each dataset. The benchmarking code is publicly available at https://github.com/zzrs123/SpaNECT.

### 4.10 Differential feature analysis

After obtaining niche labels, one-versus-rest differential feature analysis was performed for each inferred niche using the Wilcoxon rank-sum test, as implemented in the Seurat[87] R package (v5.3.1) or the Scanpy[86] Python package (v1.9.3), to identify differentially enriched genes, proteins, or peaks.

### 4.11 Functional enrichment analysis

For DLPFC, Gene Ontology (GO) enrichment analysis [74] was performed on niche-enriched genes using the clusterProfiler [94] R package (v4.19.2). For mouse brain and tonsil analyses, niche-specific pathway enrichment was performed using preranked gene set enrichment analysis [73] implemented in the gseapy [62] Python package (v1.1.8), where genes were ranked for each niche against the remaining niches based on Wilcoxon test statistics before enrichment analysis.

### 4.12 Cell-type composition and enrichment analysis

Using the inferred cell-type proportion matrix *C*, we quantified cell-type composition within each inferred niche. For niche composition analysis, cell-type proportions were aggregated across cells within each niche to obtain the mean fraction of each cell type. For cell-type enrichment analysis, an enrichment score for each cell type in each niche was defined as − log_10_(*P*), where *P* was computed using a right-tailed hypergeometric test based on four quantities: the number of cells associated with the given cell type in the niche, the total number of cells in the niche, the number of cells associated with that cell type in the full spatial map, and the total number of cells in the spatial map.

## 5 Data availability

All datasets analyzed in this study are publicly available from their original sources. Detailed information on spatial dataset sources, accession links, and modality descriptions is provided in Supplementary Table 1, and the single-cell reference datasets used for cell-type information construction are summarized in Supplementary Table 2. The processed datasets used in this study are available at Zenodo under accession code [https://doi.org/10.5281/zenodo.20064886]. Source data are provided with this paper.

## Acknowledgements

We thank all of the contributors of the open-source datasets and freely available tools used in this study. We also appreciate the suggestive comments of reviewers. This work is supported by the National Natural Science Foundation of China [62433016].

## 8 Additional information

Supporting information is available online.

## References

[1] Lázár, E., Lundeberg, J.: Spatial architecture of development and disease. Nature Reviews Genetics 27(2), 118–136 (2026)

[2] Farah, E.N., Hu, R.K., Kern, C., Zhang, Q., Lu, T.-Y., Ma, Q., Tran, S., Zhang, B., Carlin, D., Monell, A., et al.: Spatially organized cellular communities form the developing human heart. Nature 627(8005), 854–864 (2024)

[3] Vannan, A., Lyu, R., Williams, A.L., Negretti, N.M., Mee, E.D., Hirsh, J., Hirsh, S., Hadad, N., Nichols, D.S., Calvi, C.L., et al.: Spatial transcriptomics identifies molecular niche dysregulation associated with distal lung remodeling in pulmonary fibrosis. Nature genetics 57(3), 647–658 (2025)

[4] Lubeck, E., Coskun, A.F., Zhiyentayev, T., Ahmad, M., Cai, L.: Single-cell in situ rna profiling by sequential hybridization. Nature methods 11(4), 360–361 (2014)

[5] Chen, K., Boettiger, A., Moffitt, J., Wang, S., Zhuang, X.: RNA imaging. Spatially resolved, highly multiplexed RNA profiling in single cells. Science 348, aaa6090 (2015)

[6] He, S., Bhatt, R., Brown, C., Brown, E.A., Buhr, D.L., Chantranuvatana, K., Danaher, P., Dunaway, D., Garrison, R.G., Geiss, G., et al.: High-plex imaging of rna and proteins at subcellular resolution in fixed tissue by spatial molecular imaging. Nature biotechnology 40(12), 1794–1806 (2022)

[7] Zeng, H., Huang, J., Zhou, H., Meilandt, W.J., Dejanovic, B., Zhou, Y., Bohlen, C.J., Lee, S.-H., Ren, J., Liu, A., et al.: Integrative in situ mapping of single-cell transcriptional states and tissue histopathology in a mouse model of alzheimer’s disease. Nature neuroscience 26(3), 430–446 (2023)

[8] Stickels, R.R., Murray, E., Kumar, P., Li, J., Marshall, J.L., Di Bella, D.J., Arlotta, P., Macosko, E.Z., Chen, F.: Highly sensitive spatial transcriptomics at near-cellular resolution with slide-seqv2. Nature biotechnology 39(3), 313–319 (2021)

[9] Chen, A., Liao, S., Cheng, M., Ma, K., Wu, L., Lai, Y., Qiu, X., Yang, J., Xu, J., Hao, S., et al.: Spatiotemporal transcriptomic atlas of mouse organogenesis using dna nanoball-patterned arrays. Cell 185(10), 1777–1792 (2022)

[10] Liu, Y., Yang, M., Deng, Y., Su, G., Enninful, A., Guo, C.C., Tebaldi, T., Zhang, D., Kim, D., Bai, Z., et al.: High-spatial-resolution multi-omics sequencing via deterministic barcoding in tissue. Cell 183(6), 1665–1681 (2020)

[11] Fu, X., Sun, L., Dong, R., Chen, J.Y., Silakit, R., Condon, L.F., Lin, Y., Lin, S., Palmiter, R.D., Gu, L.: Polony gels enable amplifiable dna stamping and spatial transcriptomics of chronic pain. Cell 185(24), 4621–4633 (2022)

[12] Cho, C.-S., Xi, J., Si, Y., Park, S.-R., Hsu, J.-E., Kim, M., Jun, G., Kang, H.M., Lee, J.H.: Microscopic examination of spatial transcriptome using seq-scope. Cell 184(13), 3559–3572 (2021)

[13] Zhang, D., Deng, Y., Kukanja, P., Agirre, E., Bartosovic, M., Dong, M., Ma, C., Ma, S., Su, G., Bao, S., et al.: Spatial epigenome–transcriptome co-profiling of mammalian tissues. Nature 616(7955), 113–122 (2023)

[14] Jiang, F., Zhou, X., Qian, Y., Zhu, M., Wang, L., Li, Z., Shen, Q., Wang, M., Qu, F., Cui, G., et al.: Simultaneous profiling of spatial gene expression and chromatin accessibility during mouse brain development. Nature Methods 20(7), 1048–1057 (2023)

[15] Liu, Y., DiStasio, M., Su, G., Asashima, H., Enninful, A., Qin, X., Deng, Y., Nam, J., Gao, F., Bordignon, P., et al.: High-plex protein and whole transcriptome co-mapping at cellular resolution with spatial cite-seq. Nature Biotechnology 41(10), 1405–1409 (2023)

[16] Zhang, D., Rubio Rodríguez-Kirby, L.A., Lin, Y., Wang, W., Song, M., Wang, L., Wang, L., Kanatani, S., Jimenez-Beristain, T., Dang, Y., et al.: Spatial dynamics of brain development and neuroinflammation. Nature 647(8088), 213–227 (2025)

[17] Liu, L., Chen, A., Li, Y., Mulder, J., Heyn, H., Xu, X.: Spatiotemporal omics for biology and medicine. Cell 187(17), 4488–4519 (2024)

[18] Yuan, Z., Zhao, F., Lin, S., Zhao, Y., Yao, J., Cui, Y., Zhang, X.-Y., Zhao, Y.: Benchmarking spatial clustering methods with spatially resolved transcriptomics data. Nature Methods 21(4), 712–722 (2024)

[19] Zhao, E., Stone, M.R., Ren, X., Guenthoer, J., Smythe, K.S., Pulliam, T., Williams, S.R., Uytingco, C.R., Taylor, S.E., Nghiem, P., et al.: Spatial transcriptomics at subspot resolution with bayesspace. Nature biotechnology 39(11), 1375–1384 (2021)

[20] Dong, K., Zhang, S.: Deciphering spatial domains from spatially resolved transcriptomics with an adaptive graph attention auto-encoder. Nature communications 13(1), 1739 (2022)

[21] Xu, H., Fu, H., Long, Y., Ang, K.S., Sethi, R., Chong, K., Li, M., Uddamvathanak, R., Lee, H.K., Ling, J., et al.: Unsupervised spatially embedded deep representation of spatial transcriptomics. Genome Medicine 16(1), 12 (2024)

[22] Yuan, Z., Li, Y., Shi, M., Yang, F., Gao, J., Yao, J., Zhang, M.Q.: Sotip is a versatile method for microenvironment modeling with spatial omics data. Nature communications 13(1), 7330 (2022)

[23] Varrone, M., Tavernari, D., Santamaria-Martínez, A., Walsh, L.A., Ciriello, G.: Cellcharter reveals spatial cell niches associated with tissue remodeling and cell plasticity. Nature genetics 56(1), 74–84 (2024)

[24] Qian, J., Shao, X., Bao, H., Fang, Y., Guo, W., Li, C., Li, A., Hua, H., Fan, X.: Identification and characterization of cell niches in tissue from spatial omics data at single-cell resolution. Nature Communications 16(1), 1693 (2025)

[25] Long, Y., Ang, K.S., Li, M., Chong, K.L.K., Sethi, R., Zhong, C., Xu, H., Ong, Z., Sachaphibulkij, K., Chen, A., et al.: Spatially informed clustering, integration, and deconvolution of spatial transcriptomics with graphst. Nature communications 14(1), 1155 (2023)

[26] Yu, Z., Yang, Y., Chen, X., Wong, K.-C., Zhang, Z., Zhao, Y., Li, X.: Accurate spatial heterogeneity dissection and gene regulation interpretation for spatial transcriptomics using dual graph contrastive learning. Advanced Science 12(3), 2410081 (2025)

[27] Hu, Y., Rong, J., Xu, Y., Xie, R., Peng, J., Gao, L., Tan, K.: Unsupervised and supervised discovery of tissue cellular neighborhoods from cell phenotypes. Nature Methods 21(2), 267–278 (2024)

[28] Pham, D., Tan, X., Balderson, B., Xu, J., Grice, L.F., Yoon, S., Willis, E.F., Tran, M., Lam, P.Y., Raghubar, A., et al.: Robust mapping of spatiotemporal trajectories and cell–cell interactions in healthy and diseased tissues. Nature communications 14(1), 7739 (2023)

[29] Hu, J., Li, X., Coleman, K., Schroeder, A., Ma, N., Irwin, D.J., Lee, E.B., Shinohara, R.T., Li, M.: Spagcn: Integrating gene expression, spatial location and histology to identify spatial domains and spatially variable genes by graph convolutional network. Nature methods 18(11), 1342–1351 (2021)

[30] Xu, C., Jin, X., Wei, S., Wang, P., Luo, M., Xu, Z., Yang, W., Cai, Y., Xiao, L., Lin, X., et al.: Deepst: identifying spatial domains in spatial transcriptomics by deep learning. Nucleic acids research 50(22), 131–131 (2022)

[31] Yang, Y., Cui, Y., Zeng, X., Zhang, Y., Loza, M., Park, S.-J., Nakai, K.: Staig: Spatial transcriptomics analysis via image-aided graph contrastive learning for domain exploration and alignment-free integration. Nature Communications 16(1), 1067 (2025)

[32] Vandereyken, K., Sifrim, A., Thienpont, B., Voet, T.: Methods and applications for single-cell and spatial multi-omics. Nature Reviews Genetics 24(8), 494–515 (2023)

[33] Coleman, K., Schroeder, A., Loth, M., Zhang, D., Park, J.H., Sung, J.-Y., Blank, N., Cowan, A.J., Qian, X., Chen, J., et al.: Resolving tissue complexity by multimodal spatial omics modeling with miso. Nature methods 22(3), 530–538 (2025)

[34] Li, Z., Qu, S., Liang, H., Tang, R., Zhang, X., Lu, F., Yang, J., Gan, Z., Gao, S., Zhang, Y., et al.: Integrative deep learning of spatial multi-omics with switch. Nature Computational Science, 1–13 (2025)

[35] Miao, J., Li, J., Xin, J., Tu, J., Ge, M., Qi, J., Zhou, X., Zhu, Y., Yang, C., Lin, Z.: Multigate: integrative analysis and regulatory inference in spatial multiomics data via graph representation learning. Nature Communications 16(1), 9403 (2025)

[36] Long, Y., Ang, K.S., Sethi, R., Liao, S., Heng, Y., Olst, L., Ye, S., Zhong, C., Xu, H., Zhang, D., et al.: Deciphering spatial domains from spatial multi-omics with spatialglue. Nature Methods 21(9), 1658–1667 (2024)

[37] Zhou, Y., Xiao, X., Dong, L., Tang, C., Xiao, G., Xu, L.: Cooperative integration of spatially resolved multi-omics data with cosmos. Nature communications 16(1), 27 (2025)

[38] Liu, X., Peng, T., Xu, M., Lin, S., Hu, B., Chu, T., Liu, B., Xu, Y., Ding, W., Li, L., et al.: Spatial multi-omics: deciphering technological landscape of integration of multi-omics and its applications. Journal of Hematology & Oncology 17(1), 72 (2024)

[39] Chelebian, E., Avenel, C., Wählby, C.: Combining spatial transcriptomics with tissue morphology. Nature Communications 16(1), 4452 (2025)

[40] Maynard, K.R., Collado-Torres, L., Weber, L.M., Uytingco, C., Barry, B.K., Williams, S.R., Catallini, J.L., Tran, M.N., Besich, Z., Tippani, M., et al.: Transcriptome-scale spatial gene expression in the human dorsolateral prefrontal cortex. Nature neuroscience 24(3), 425–436 (2021)

[41] Chen, S., Chang, Y., Li, L., Acosta, D., Li, Y., Guo, Q., Wang, C., Turkes, E., Morrison, C., Julian, D., et al.: Spatially resolved transcriptomics reveals genes associated with the vulnerability of middle temporal gyrus in alzheimer’s disease. Acta neuropathologica communications 10(1), 188 (2022)

[42] Yan, X., Ang, K.S., Olst, L., Edwards, A., Watson, T., Zheng, R., Fan, R., Li, M., Gate, D., Chen, J.: Mosaic integration of spatial multi-omics with spamosaic. BioRxiv, 2024–10 (2024)

[43] Mantri, M., Scuderi, G.J., Abedini-Nassab, R., Wang, M.F., McKellar, D., Shi, H., Grodner, B., Butcher, J.T., De Vlaminck, I.: Spatiotemporal single-cell rna sequencing of developing chicken hearts identifies interplay between cellular differentiation and morphogenesis. Nature communications 12(1), 1771 (2021)

[44] Tran, M.N., Maynard, K.R., Spangler, A., Huuki, L.A., Montgomery, K.D., Sadashivaiah, V., Tippani, M., Barry, B.K., Hancock, D.B., Hicks, S.C., et al.: Single-nucleus transcriptome analysis reveals cell-type-specific molecular signatures across reward circuitry in the human brain. Neuron 109(19), 3088–3103 (2021)

[45] Hodge, R.D., Bakken, T.E., Miller, J.A., Smith, K.A., Barkan, E.R., Graybuck, L.T., Close, J.L., Long, B., Johansen, N., Penn, O., et al.: Conserved cell types with divergent features in human versus mouse cortex. Nature 573(7772), 61–68 (2019)

[46] Rajkowska, G., Mahajan, G., Maciag, D., Sathyanesan, M., Iyo, A.H., Moulana, M., Kyle, P.B., Woolverton, W.L., Miguel-Hidalgo, J.J., Stockmeier, C.A., et al.: Oligodendrocyte morphometry and expression of myelin–related mrna in ventral prefrontal white matter in major depressive disorder. Journal of psychiatric research 65, 53–62 (2015)

[47] Hercher, C., Chopra, V., Beasley, C.L.: Evidence for morphological alterations in prefrontal white matter glia in schizophrenia and bipolar disorder. Journal of Psychiatry and Neuroscience 39(6), 376–385 (2014)

[48] Shirzadi, Z., Schultz, S.A., Yau, W.-Y.W., Joseph-Mathurin, N., Fitzpatrick, C.D., Levin, R., Kantarci, K., Preboske, G.M., Jack Jr, C.R., Farlow, M.R., et al.: Etiology of white matter hyperintensities in autosomal dominant and sporadic alzheimer disease. JAMA neurology 80(12), 1353–1363 (2023)

[49] Nasrabady, S.E., Rizvi, B., Goldman, J.E., Brickman, A.M.: White matter changes in alzheimer’s disease: a focus on myelin and oligodendrocytes. Acta neuropathologica communications 6(1), 22 (2018)

[50] Burgos-Ramos, E., Hervás-Aguilar, A., Aguado-Llera, D., Puebla-Jiménez, L., Hernández-Pinto, A., Barrios, V., Arilla-Ferreiro, E.: Somatostatin and alzheimer’s disease. Molecular and cellular endocrinology 286(1-2), 104–111 (2008)

[51] Tayran, H., Yilmaz, E., Bhattarai, P., Min, Y., Wang, X., Ma, Y., Wang, N., Jeong, I., Nelson, N., Kassara, N., et al.: Abca7-dependent induction of neuropeptide y is required for synaptic resilience in alzheimer’s disease through bdnf/ngfr signaling. Cell genomics 4(9) (2024)

[52] Xu, Y., Shao, N., Zhi, F., Chen, R., Yang, Y., Li, J., Xia, Y., Peng, Y.: Delta-opioid receptor signaling alleviates neuropathology and cognitive impairment in the mouse model of alzheimer’s disease by regulating microglia homeostasis and inhibiting hmgb1 pathway. Alzheimer’s Research & Therapy 17(1), 35 (2025)

[53] Gao, C., Jiang, J., Tan, Y., Chen, S.: Microglia in neurodegenerative diseases: mechanism and potential therapeutic targets. Signal transduction and targeted therapy 8(1), 359 (2023)

[54] Ceci, C., Lacal, P.M., Barbaccia, M.L., Mercuri, N.B., Graziani, G., Ledonne, A.: The vegfs/vegfrs system in alzheimer’s and parkinson’s diseases: Pathophysiological roles and therapeutic implications. Pharmacological Research 201, 107101 (2024)

[55] Wang, H., Lv, J.-J., Zhao, Y., Wei, H.-L., Zhang, T.-J., Yang, H.-J., Chen, Z.-N., Jiang, J.-L.: Endothelial genetic deletion of cd147 induces changes in the dual function of the blood-brain barrier and is implicated in alzheimer’s disease. CNS neuroscience & therapeutics 27(9), 1048–1063 (2021)

[56] Pan, P., Zhao, H., Zhang, X., Li, Q., Qu, J., Zuo, S., Yang, F., Liang, G., Zhang, J.H., Liu, X., et al.: Cyclophilin a signaling induces pericyte-associated blood-brain barrier disruption after subarachnoid hemorrhage. Journal of neuroinflammation 17(1), 16 (2020)

[57] Bell, R.D., Winkler, E.A., Singh, I., Sagare, A.P., Deane, R., Wu, Z., Holtzman, D.M., Betsholtz, C., Armulik, A., Sallstrom, J., et al.: Apolipoprotein e controls cerebrovascular integrity via cyclophilin a. Nature 485(7399), 512–516 (2012)

[58] Rahimian, R., Perlman, K., Canonne, C., Mechawar, N.: Targeting microglia– oligodendrocyte crosstalk in neurodegenerative and psychiatric disorders. Drug Discovery Today 27(9), 2562–2573 (2022)

[59] La Manno, G., Siletti, K., Furlan, A., Gyllborg, D., Vinsland, E., Mossi Albiach, A., Mattsson Langseth, C., Khven, I., Lederer, A.R., Dratva, L.M., et al.: Molecular architecture of the developing mouse brain. Nature 596(7870), 92–96 (2021)

[60] Galceran, J., Miyashita-Lin, E.M., Devaney, E., Rubenstein, J.L., Grosschedl, R.: Hippocampus development and generation of dentate gyrus granule cells is regulated by lef1. Development 127(3), 469–482 (2000)

[61] Sagai, T., Amano, T., Maeno, A., Ajima, R., Shiroishi, T.: Shh signaling mediated by a prechordal and brain enhancer controls forebrain organization. Proceedings of the National Academy of Sciences 116(47), 23636–23642 (2019)

[62] Fang, Z., Liu, X., Peltz, G.: Gseapy: a comprehensive package for performing gene set enrichment analysis in python. Bioinformatics 39(1), 757 (2023)

[63] Pavert, S.A., Mebius, R.E.: New insights into the development of lymphoid tissues. Nature Reviews Immunology 10(9), 664–674 (2010)

[64] Massoni-Badosa, R., Aguilar-Fernández, S., Nieto, J.C., Soler-Vila, P., Elosua-Bayes, M., Marchese, D., Kulis, M., Vilas-Zornoza, A., Bühler, M.M., Rashmi, S., et al.: An atlas of cells in the human tonsil. Immunity 57(2), 379–399 (2024)

[65] Cyster, J.G., Allen, C.D.: B cell responses: cell interaction dynamics and decisions. Cell 177(3), 524–540 (2019)

[66] Mueller, S.N., Germain, R.N.: Stromal cell contributions to the homeostasis and functionality of the immune system. Nature Reviews Immunology 9(9), 618–629 (2009)

[67] Perry, M., Whyte, A.: Immunology of the tonsils. Immunology today 19(9), 414–421 (1998)

[68] Chen, S., Zhu, B., Huang, S., Hickey, J.W., Lin, K.Z., Snyder, M., Greenleaf, W.J., Nolan, G.P., Zhang, N.R., Ma, Z.: Integration of spatial and single-cell data across modalities with weakly linked features. Nature Biotechnology 42(7), 1096–1106 (2024)

[69] Schebesta, M., Heavey, B., Busslinger, M.: Transcriptional control of b-cell development. Current opinion in immunology 14(2), 216–223 (2002)

[70] Förster, R., Schubel, A., Breitfeld, D., Kremmer, E., Renner-Müller, I., Wolf, E., Lipp, M.: Ccr7 coordinates the primary immune response by establishing functional microenvironments in secondary lymphoid organs. Cell 99(1), 23–33 (1999)

[71] Kapellos, T.S., Bonaguro, L., Gemünd, I., Reusch, N., Saglam, A., Hinkley, E.R., Schultze, J.L.: Human monocyte subsets and phenotypes in major chronic inflammatory diseases. Frontiers in immunology 10, 2035 (2019)

[72] Havenar-Daughton, C., Lindqvist, M., Heit, A., Wu, J.E., Reiss, S.M., Kendric, K., Bélanger, S., Kasturi, S.P., Landais, E., Akondy, R.S., et al.: Cxcl13 is a plasma biomarker of germinal center activity. Proceedings of the National Academy of Sciences 113(10), 2702–2707 (2016)

[73] Subramanian, A., Tamayo, P., Mootha, V.K., Mukherjee, S., Ebert, B.L., Gillette, M.A., Paulovich, A., Pomeroy, S.L., Golub, T.R., Lander, E.S., et al.: Gene set enrichment analysis: a knowledge-based approach for interpreting genome-wide expression profiles. Proceedings of the national academy of sciences 102(43), 15545–15550 (2005)

[74] The gene ontology resource: enriching a gold mine. Nucleic acids research 49(D1), 325–334 (2021)

[75] Mesin, L., Ersching, J., Victora, G.D.: Germinal center b cell dynamics. Immunity 45(3), 471–482 (2016)

[76] Meilhac, S.M., Buckingham, M.E.: The deployment of cell lineages that form the mammalian heart. Nature Reviews Cardiology 15(11), 705–724 (2018)

[77] MacGrogan, D., Münch, J., Pompa, J.L.: Notch and interacting signalling pathways in cardiac development, disease, and regeneration. Nature Reviews Cardiology 15(11), 685–704 (2018)

[78] Lupu, I.-E., De Val, S., Smart, N.: Coronary vessel formation in development and disease: mechanisms and insights for therapy. Nature Reviews Cardiology 17(12), 790–806 (2020)

[79] Butcher, J.T., Markwald, R.R.: Valvulogenesis: the moving target. Philosophical Transactions of the Royal Society B: Biological Sciences 362(1484), 1489–1503 (2007)

[80] Grego-Bessa, J., Luna-Zurita, L., Monte, G., Bolós, V., Melgar, P., Arandilla, A., Garratt, A.N., Zang, H., Mukouyama, Y.-s., Chen, H., et al.: Notch signaling is essential for ventricular chamber development. Developmental cell 12(3), 415–429 (2007)

[81] Ryan, A.K., Blumberg, B., Rodriguez-Esteban, C., Yonei-Tamura, S., Tamura, K., Tsukui, T., De La Peña, J., Sabbagh, W., Greenwald, J., Choe, S., et al.: Pitx2 determines left–right asymmetry of internal organs in vertebrates. Nature 394(6693), 545–551 (1998)

[82] Lin, C.R., Kioussi, C., O’Connell, S., Briata, P., Szeto, D., Liu, F., Izpisuá-Belmonte, J.C., Rosenfeld, M.G.: Pitx2 regulates lung asymmetry, cardiac positioning and pituitary and tooth morphogenesis. Nature 401(6750), 279–282 (1999)

[83] Koshiba-Takeuchi, K., Mori, A.D., Kaynak, B.L., Cebra-Thomas, J., Sukonnik, T., Georges, R.O., Latham, S., Beck, L., Henkelman, R.M., Black, B.L., et al.: Reptilian heart development and the molecular basis of cardiac chamber evolution. Nature 461(7260), 95–98 (2009)

[84] Wu, E., Bieniosek, M., Wu, Z., Thakkar, N., Charville, G.W., Makky, A., Schürch, C.M., Huyghe, J.R., Peters, U., Li, C.I., et al.: Rosie: Ai generation of multiplex immunofluorescence staining from histopathology images. Nature Communications 16(1), 7633 (2025)

[85] Cao, Y., Zhao, X., Tang, S., Jiang, Q., Li, S., Li, S., Chen, S.: scbutterfly: a versatile single-cell cross-modality translation method via dual-aligned variational autoencoders. Nature Communications 15(1), 2973 (2024)

[86] Wolf, F.A., Angerer, P., Theis, F.J.: Scanpy: large-scale single-cell gene expression data analysis. Genome biology 19(1), 15 (2018)

[87] Stuart, T., Butler, A., Hoffman, P., Hafemeister, C., Papalexi, E., Mauck, W.M., Hao, Y., Stoeckius, M., Smibert, P., Satija, R.: Comprehensive integration of single-cell data. cell 177(7), 1888–1902 (2019)

[88] Grill, J.-B., Strub, F., Altché, F., Tallec, C., Richemond, P., Buchatskaya, E., Doersch, C., Avila Pires, B., Guo, Z., Gheshlaghi Azar, M., et al.: Bootstrap your own latent-a new approach to self-supervised learning. Advances in neural information processing systems 33, 21271–21284 (2020)

[89] Elosua-Bayes, M., Nieto, P., Mereu, E., Gut, I., Heyn, H.: Spotlight: seeded nmf regression to deconvolute spatial transcriptomics spots with single-cell transcriptomes. Nucleic acids research 49(9), 50–50 (2021)

[90] Ma, Y., Zhou, X.: Spatially informed cell-type deconvolution for spatial transcriptomics. Nature biotechnology 40(9), 1349–1359 (2022)

[91] Kinga, D., Adam, J.B., et al.: A method for stochastic optimization. In: International Conference on Learning Representations (ICLR), vol. 5 (2015). California;

[92] Hartigan, J.A., Wong, M.A.: Algorithm as 136: A k-means clustering algorithm. Journal of the royal statistical society. series c (applied statistics) 28(1), 100–108 (1979)

[93] Scrucca, L., Fop, M., Murphy, T.B., Raftery, A.E.: mclust 5: clustering, classification and density estimation using gaussian finite mixture models. The R journal 8(1), 289 (2016)

[94] Yu, G., Wang, L.-G., Han, Y., He, Q.-Y.: clusterprofiler: an r package for comparing biological themes among gene clusters. Omics: a journal of integrative biology 16(5), 284–287 (2012)

